# Positional dynamics and glycosomal recruitment of developmental regulators during trypanosome differentiation

**DOI:** 10.1101/603258

**Authors:** Balázs Szöőr, Dorina V. Simon, Federico Rojas, Julie Young, Derrick R. Robinson, Timothy Krüger, Markus Engstler, Keith R. Matthews

## Abstract

Glycosomes are peroxisome-related organelles that compartmentalise the glycolytic enzymes in kinetoplastid parasites. These organelles are developmentally regulated in their number and composition, allowing metabolic adaptation to the parasite’s needs in the blood of mammalian hosts or within their arthropod vector. A protein phosphatase cascade regulates differentiation between parasite developmental forms, comprising a tyrosine phosphatase, TbPTP1, that dephosphorylates and inhibits a serine threonine phosphatase TbPIP39 that promotes differentiation. When TbPTP1 is inactivated, TbPIP39 is activated and during differentiation becomes located in glycosomes. Here we have tracked TbPIP39 recruitment to glycosomes during differentiation from bloodstream stumpy forms to procyclic forms. Detailed microscopy and live cell imaging during the synchronous transition between life cycle stages revealed that in stumpy forms, TbPIP39 is located at a periflagellar pocket site closely associated with TbVAP, that defines the flagellar pocket endoplasmic reticulum. TbPTP1 is also located at the same site in stumpy forms, as is REG9.1, a regulator of stumpy-enriched mRNAs. This site provides a molecular node for the interaction between TbPTP1 and TbPIP39. Within 30 minutes of the initiation of differentiation TbPIP39 relocates to glycosomes whereas TbPTP1 disperses to the cytosol. Overall, the study identifies a ‘stumpy regulatory nexus’ (STuRN) that co-ordinates the molecular components of life cycle signalling and glycosomal development during transmission of *Trypanosoma brucei*.

**Importance:** African trypanosomes are parasites of sub-Saharan Africa responsible for both human and animal disease. The parasites are transmitted by tsetse flies and completion of their life cycle involves progression through several development steps. The initiation of differentiation between blood and tsetse forms is signalled by a phosphatase cascade, ultimately trafficked into peroxisome-related organelles called glycosomes that are unique to this group of organisms. Glycosomes undergo substantial remodelling of their composition and function during the differentiation step but how this is regulated is not understood. Here we identify a cytological site where the signalling molecules controlling differentiation converge before the dispersal of one of them into glycosomes. This coincides with a specialised ER site that may contribute to glycosome developmental biogenesis or regeneration. In combination, the study provides the first insight into the spatial co-ordination of signalling pathway components in trypanosomes as they undergo cell-type differentiation.

## Introduction

The dynamic regulation of organelle biogenesis or composition often involves intimate contact between the endoplasmic reticulum and the membrane of the target organelle (1). This enables the maturation and modification of organellar protein content, influencing mitochondrial, Golgi or peroxisomal components, whereas inter-organellar contacts can also contribute to signalling events within cells, bringing regulatory molecules into proximity, or trafficking them for degradation (2). The protein contacts at the interface between organelles are often diverse and characteristic of each organellar type, predominantly interacting with vesicle associated membrane protein associated proteins (VAPs) on the ER membrane.

In addition to the conventional organelles typical of eukaryotic cells, evolutionarily divergent kinetoplastid parasites are characterised by their possession of glycosomes, specialist organelles that harbour the enzymes of glycolysis (3). Although unique in their compartmentation of glycolytic enzymes, glycosomes are related to peroxisomes, sharing with those organelles a similar (though divergent) machinery for import, insertion of membrane proteins (PEX16, PEX19) and peroxisome proliferation, as well as their capacity for either lipid, purine and pyrimidine biosynthesis (4). Glycosomes are also dynamic in composition and number in response to the metabolic demands of the parasite, their synthesis and turnover involving similar biogenesis and degradation mechanisms to those of the peroxisomes of yeast and mammalian cells. This capacity for biosynthesis and turnover enables peroxisomes and glycosomes to exploit different nutrient conditions or adapt to different developmental forms.

Kinetoplastid parasites comprise pathogens of mammals that are frequently transmitted by arthropod vectors. Among the best characterised and tractable are the African trypanosomes, *Trypanosoma brucei*. These parasites live extracellularly in the bloodstream and tissues of mammalian hosts, where they cause human sleeping sickness and the livestock disease nagana (5, 6). Trypanosomes are spread by blood feeding tsetse flies, the passage between the blood and the insect gut involving a switch from glucose based energy metabolism to one reliant on amino acids (7). Pivotal to the successful colonisation of the tsetse fly are so called ‘stumpy forms’, quiescent bloodstream forms that show several adaptations for survival upon uptake by tsetse flies (8), including partial elaboration of their mitochondrion in preparation for the switch from glucose-dependent energy generation *via* glycolysis (9–12). Stumpy forms arise from proliferative slender forms in the bloodstream in a quorum sensing response dependent upon parasite density (13). This results in the accumulation of uniform populations of stumpy forms that are cell cycle arrested in G1/G0 and sensitised for differentiation when taken up in a tsetse fly blood meal (14), this culminating in the production of a population of differentiated procyclic forms that colonise the tsetse midgut. The same transition can also be enacted *in vitro* by exposing stumpy forms to reduced temperature and *cis*-aconitate/citrate, this generating a highly synchronised differentiation model allowing cytological events to be readily tracked and quantitated in the population (15).

The signalling events that stimulate the differentiation of stumpy forms to procyclic forms are quite well characterised. Thus, stumpy forms are held poised for differentiation by the action of a negative regulator of differentiation, TbPTP1, a tyrosine specific phosphatase (16, 17). A substrate of TbPTP1 is the DxDXT/V class serine threonine phosphatase TbPIP39, which is dephosphorylated on tyrosine 278 by TbPTP1, this interaction reducing the activity of TbPIP39 and so preventing differentiation (18). When exposed to reduced temperature, as would occur during a tsetse bloodmeal (19), blood citrate is transported by ‘PAD’ proteins whose expression is elevated on stumpy forms at 20°C (20). When exposed to citrate/*cis*-aconitate, TbPTP1 is inactivated and TbPIP39 becomes phosphorylated and activated, this stimulating differentiation of the parasites. Interestingly, sequence analysis of TbPIP39 revealed the presence of a PTS1 glycosomal localisation motif (-SRL) and this localisation was confirmed in procyclic forms by both its co-localisation with glycosomal markers (17) and its detection in glycosomal proteome analysis (21). This linked differentiation signalling in the bloodstream form parasites with glycosomal signalling during differentiation, with TbPIP39 being expressed in stumpy forms but not slender forms and being localised in glycosomes in procyclic forms.

Here we have exploited the differential expression and glycosomal location of TbPIP39 to explore the spatial positioning of differentiation signalling molecules during the transformation of stumpy forms to procyclic forms. Our results reveal the coincidence of TbPIP39 and TbPTP1 in bloodstream stumpy forms at a novel periflagellar pocket location, closely associated with a flagellar pocket ER contact site defined by TbVAP (22). This provides a novel signalling response linking environmental perception with organellar dynamics during the developmental cycle of the parasites and provides the earliest yet identified event in the initiation of trypanosome differentiation.

## Results

### TbPIP39 redistributes during synchronous differentiation

TbPIP39 is expressed in stumpy forms but not slender forms, and is maintained in established procyclic forms in glycosomes (17). To assay the recruitment of TbPIP39 to glycosomes during differentiation we analysed parasites undergoing synchronous development from stumpy to procyclic forms. Specifically, a stumpy enriched (80-90%) population was stimulated to undergo differentiation by exposure to 6mM *cis*-aconitate *in vitro*. Interestingly, the stumpy form location of TbPIP39 was distinct from that of the glycosomal marker protein glycosomal triosephosphate isomerase (gTIM). Rather than being detected in a punctate location throughout the cell typical of glycosomal staining, there was a concentration of staining close to the enlarged flagellar pocket, positioned between the nucleus and kinetoplast of the cell (Figure 1A). In contrast, procyclic forms exhibited the expected distribution of TbPIP39, where co-localisation with both glycosomal aldolase and gTIM was evident. To quantitate this location, slender, stumpy and procyclic form parasites were co-stained for TbPIP39 and gTIM, or for aldolase and gTIM and analysed by confocal microscopy, with co-localisation determined by global Pearson correlation using Volocity software (Figure 1B). This confirmed that aldolase and gTIM showed correlation of >0.8 in each life cycle stage consistent with the ubiquitous expression of these marker glycosomal proteins. In contrast, the correlation of TbPIP39 and gTIM was less than 0.15 in slender forms (where TbPIP39 is not expressed at significant levels), and less than 0.2 in stumpy forms (where TbPIP39 is expressed), whilst in procyclic forms the correlation was over 0.8 This indicated that TbPIP39 may be recruited to glycosomes during the differentiation between bloodstream and procyclic forms.

**Figure 1.**
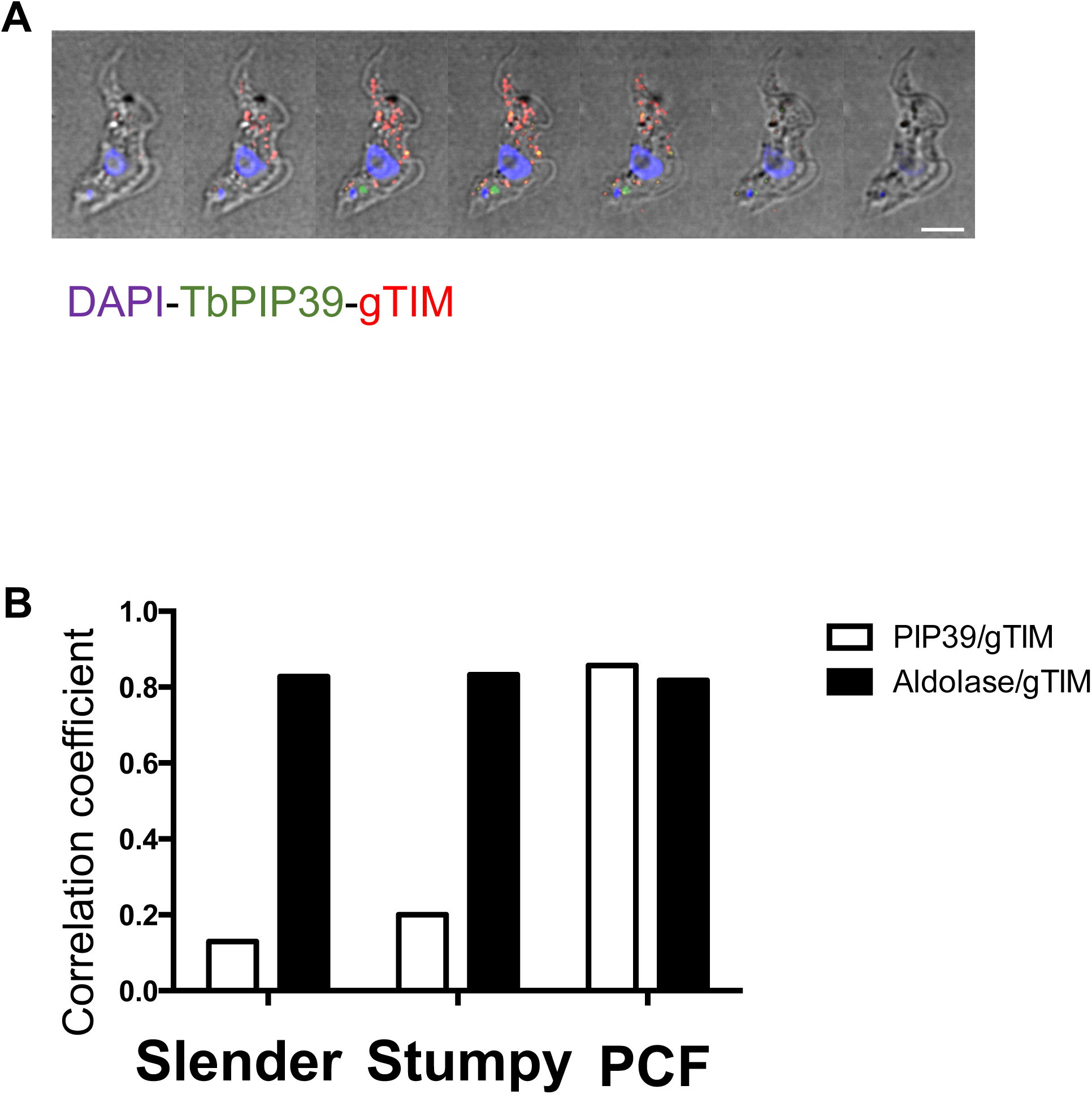
A. Serial Z stack slices through a stumpy form trypanosome cell stained to localise the differentiation regulator TbPIP39 (green) or glycosomal TIM (red). The cell nucleus and kinetoplast are shown in blue. The TbPIP39 is located close to, but slightly anterior of, the kinetoplast and is not collocated with the glycosomal marker. Bar=5µm.
B. Pearson coefficient of colocalisation between TbPIP39 and glycosomal TIM or between aldolase and glycosomal TIM in bloodstream slender and stumpy forms, or in procyclic forms. Colocalisation values were calculated using Volocity software based on captured confocal images. Threshold was set according to background of images.

To analyse the kinetics of recruitment, the distribution of TbPIP39 and gTIM were analysed at time points after exposure of stumpy form parasites to 6mM *cis*-aconitate *in vitro*. Figure 2A, B demonstrates that the periflagellar pocket staining of TbPIP39 was detectable at 0, 20 and 60 minutes following exposure to *cis*-aconitate, but was lost beyond that point. Conversely, a punctate glycosomal signal for TbPIP39 that co-localised with gTIM was detectable by 20 minutes, such that both a glycosomal and periflagellar location was present for TbPIP39 at this early time point. At 60 minutes the periflagellar staining was still detectable on some cells but the glycosomal staining was more emphasised and beyond this timepoint, the staining was mainly glycosomal.

**Figure 2.**
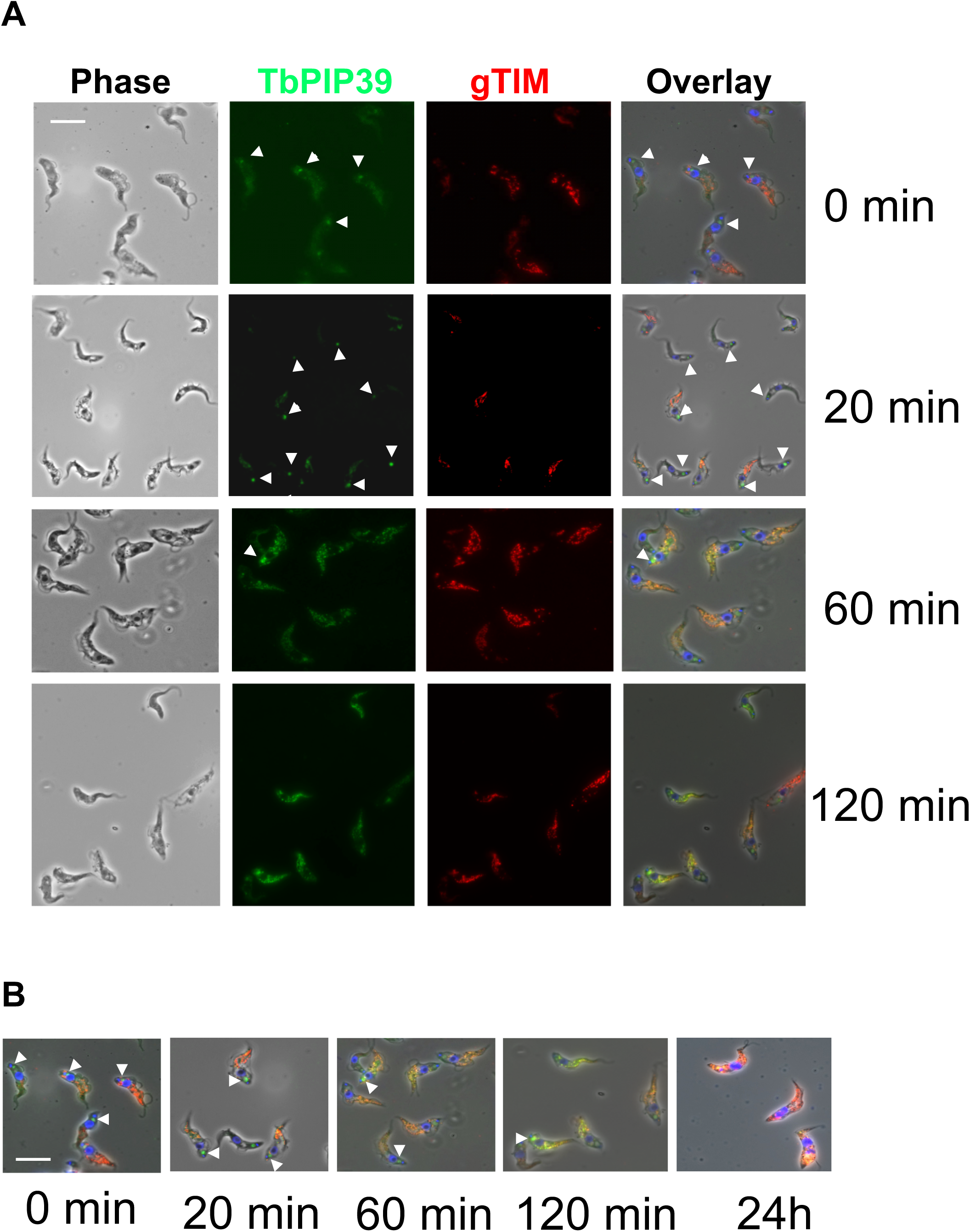
A. Representative images of trypanosomes undergoing differentiation from stumpy forms to procyclic forms. Samples were taken at the time points indicated after the initiation of differentiation by *cis*-aconitate. Cells were labelled for the location of TbPIP39 (green) or the glycosomal marker gTIM (red), with the nucleus and kinetoplast being labelled with DAPI (blue). Phase contrast images are shown on the left-hand side and merged images shown on the right. Bar= 10µm.
B. Selected fields of cells stained for TbPIP39, gTIM or DAPI at time points after the initiation of differentiation. The location of TbPIP39 proximally to the flagellar pocket region of the cell is highlighted with arrowheads. Bar=12µm.

To quantitate the redistribution of the TbPIP39 signal, 250 cells were scored at each time point after exposure to *cis*-aconitate and the parasites assayed for signal either at the periflagellar pocket region alone, at the periflagellar pocket region and glycosomes or in glycosomes alone. Figure 3A demonstrates that there was a transition during differentiation, with a predominantly glycosomal signal evident at 30 minutes and beyond in the differentiation time course. In contrast, parasites maintained *in vitro* at 27 ° C without *cis*-aconitate retained the signal close to the flagellar pocket, although there was also some glycosomal staining evident in some (20-50%) of the cells at 20 minutes and beyond. However, few (5%) cells exhibited only glycosomal staining, unlike when differentiation was stimulated with *cis*-aconitate, where >95% exhibited exclusively glycosomal location at 24h. A global Pearson correlation analysis of parasites undergoing differentiation demonstrated that the colocalization of TbPIP39 and gTIM increased from around 30% at 30 minutes to nearly 80% at 120 minutes, highlighting the rapidity of the redistribution. In contrast, in the absence of *cis*-aconitate, the correlation never exceeded 40% (Figure 3B). The correlation between aldolase and gTIM in the same analysis in contrast was consistently above 80% demonstrating the stability of the co-localisation between these glycosomal markers through the differentiation time course.

**Figure 3.**
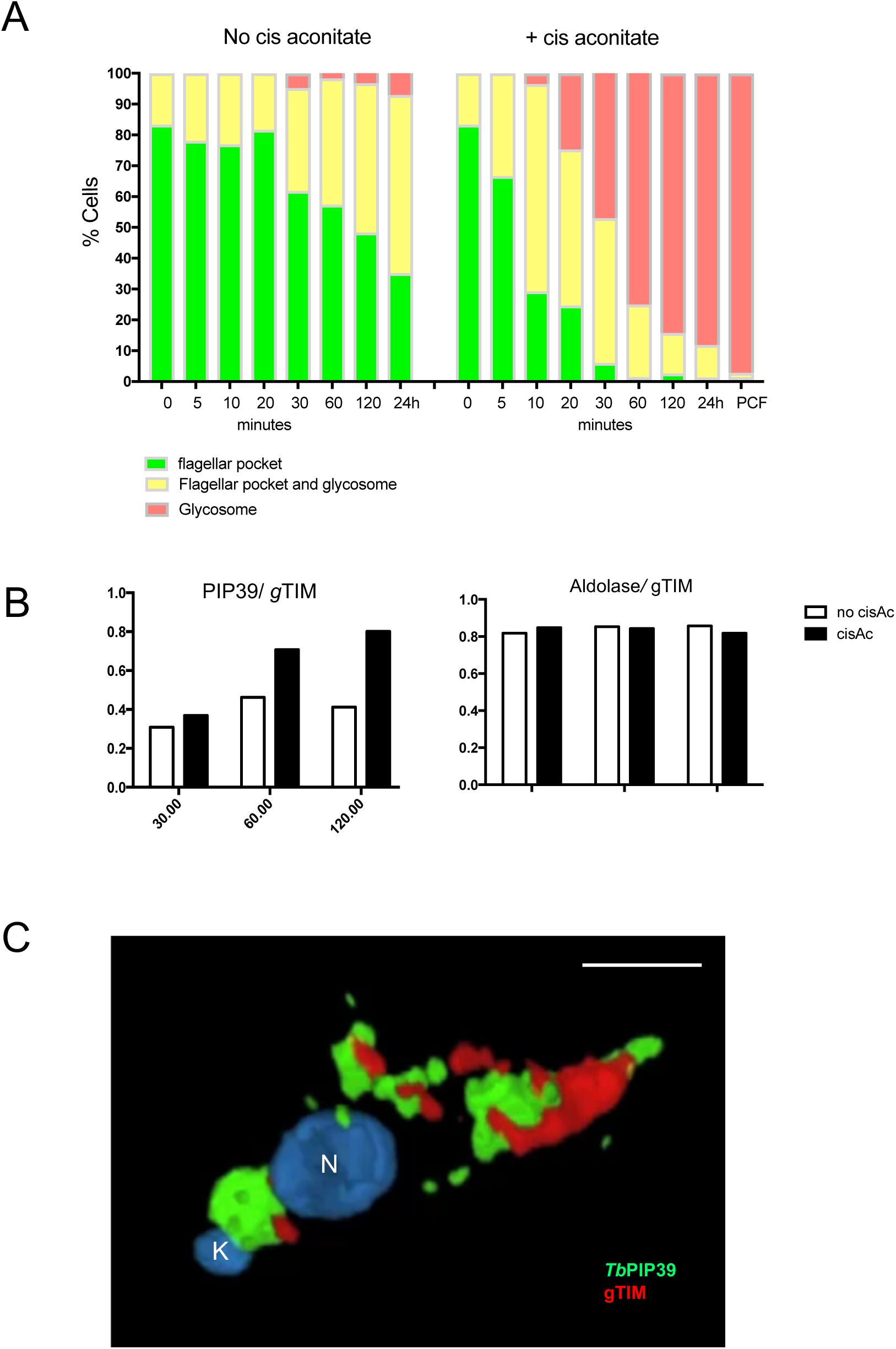
A. Quantitation of the distribution of TbPIP39 between the periflagellar pocket location only (green), at the periflagellar pocket and in glycosomes (yellow) or exclusively in glycosomes (red). At each time point and in each condition (with or without *cis*-aconitate, to initiate differentiation), 250 cells were scored.
B. Pearson coefficient of colocalisation between TbPIP39 and glycosomal TIM or between aldolase and glycosomal TIM at time points after the exposure of stumpy forms to *cis*-aconitate. Colocalisation values were calculated using Volocity software based on captured confocal images. The threshold was set according to background of images.
C. 3-D reconstruction of a cell 30 minutes after exposure to *cis*-aconitate and stained for TbPIP39 (green) and glycosomal gTIM (red). The cell nucleus (N) and kinetoplast (K) are labelled blue. TbPIP39 is concentrated around the flagellar pocket of the cell but also shows labelling at a dispersed glycosomal location more anterior in the cell, coincident with the distribution of glycosomal gTIM. Bar=5µm.

### Cytological analysis of the periflagellar staining of TbPIP39 in stumpy and differentiating cells

To examine the unusual periflagellar pocket location of TbPIP39 in stumpy forms more closely, we carried out confocal microscopy to visualise the three-dimensional distribution of TbPIP39 at 30 minutes after exposure to *cis*-aconitate. Figure 3C shows the concentration of the TbPIP39 around the flagellar pocket of the cell. As expected at 30 minutes after exposure to *cis*-aconitate, TbPIP39 was also detected in the glycosomal material anterior of the nucleus. Cells were also examined by deconvolution fluorescence microscopy after labelling for TbPIP39 and the lysosomal marker, p67 (Figure 4A). This again demonstrated the periflagellar pocket location of TbPIP39 but revealed that the signal was not evenly distributed around the flagellar pocket periphery but rather concentrated at discrete foci around the pocket. Furthermore, in cells where the flagellar pocket was collapsed during fixation, the TbPIP39 focused to a tight point supporting the distribution of the signal around, not in the flagellar pocket (Figure 4B). The site of TbPIP39 localisation was also investigated by immunoelectron microscopy (Figure 5A-E). Although signal was not abundant we were able to detect clusters of TbPIP39 labelling within the lumen of membrane bound structures and at vesicle membranes distinct from the flagellar pocket membrane and close to glycosomes in this region of the cell.

**Figure 4.**
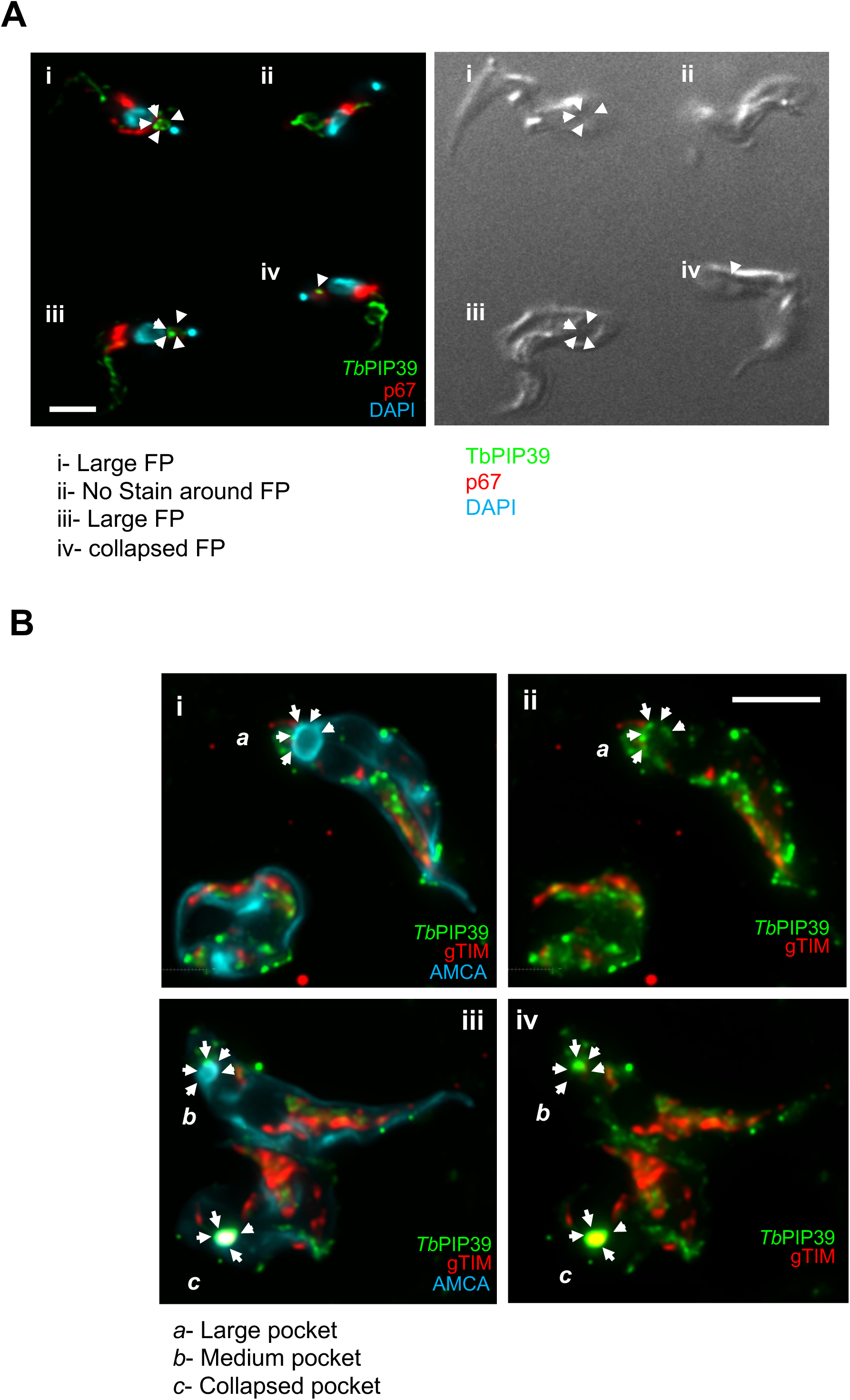
A. Co-labelling of stumpy form cells (without exposure to *cis*-aconitate) stained for TbPIP39 (green) and the lysosomal marker p67 (red). The nucleus and kinetoplast are stained in blue. The TbPIP39 labelling is distinct from the lysosome, being positioned unevenly around the flagellar pocket (cell i) or at a tight focus in cells where the flagellar pocket is collapsed (Cell iii, iv). Cell ii has no staining detected at the flagellar pocket region. Bar=5µm.
B. Co-labelling of stumpy form cells (without exposure to *cis*-aconitate) stained for TbPIP39 (green) and the glycosomal gTIM (red). The flagellar pocket and cell surface membrane are labelled blue with AMCA. Arrows indicate the distribution of the TbPIP39 signal unevenly around the flagellar pocket (Cell *a* image i, ii; Cell *b*, image iii, iv), or at a tight focus associated with a collapsed flagellar pocket (Cell *c*, image iii, iv). Bar=5µm

**Figure 5.**
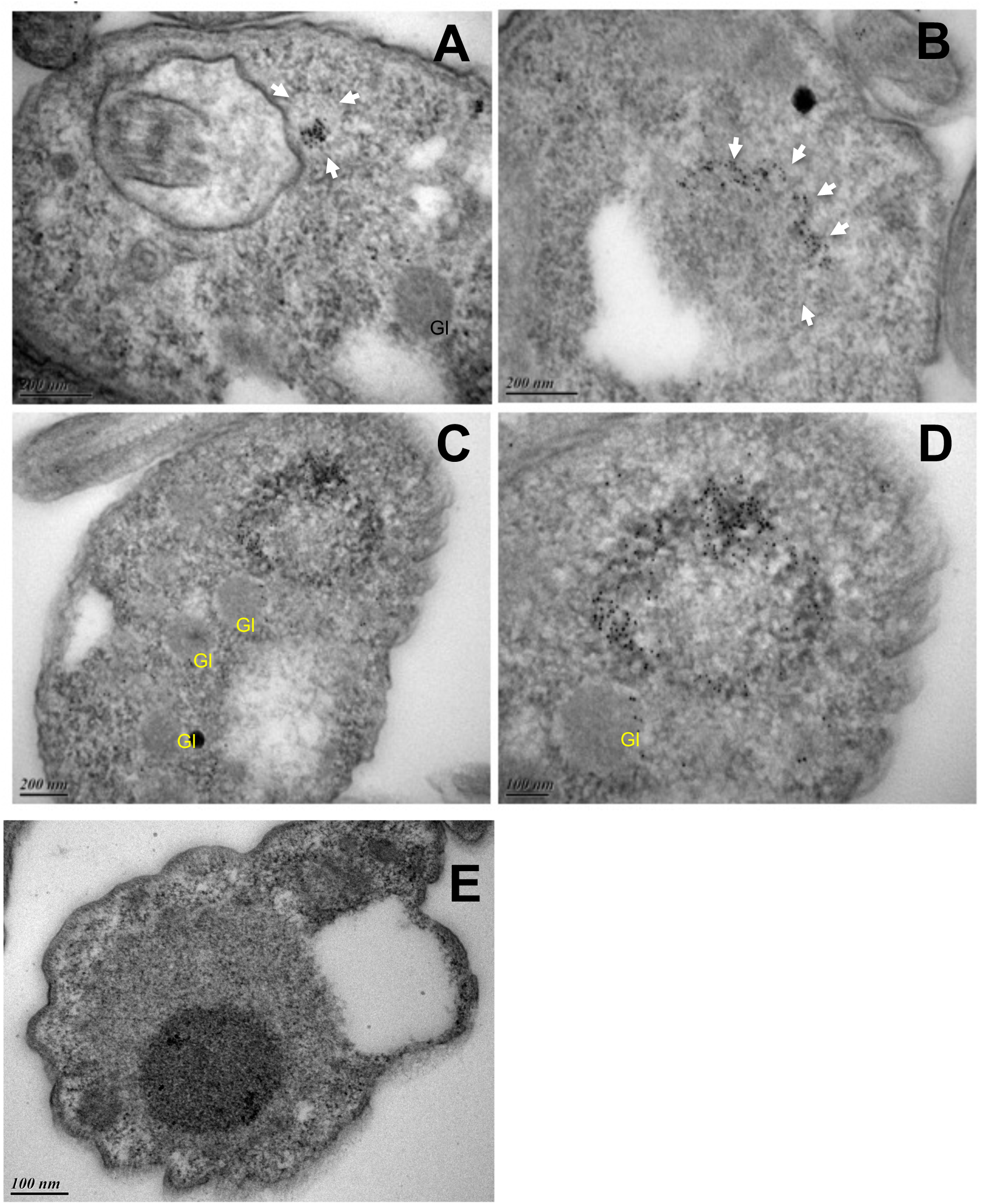
A-D Immunoelectron micrographs taken of thin sections of the flagellar pocket region of stumpy cells immunogold-labelled for TbPIP39. Arrows indicate the boundary of membrane bound vesicles containing TbPIP39 signal; glycosomes are labelled ‘Gl’.
E Immunoelectron micrographs of the flagellar pocket region of stumpy cells in the absence of primary antibody.

Finally, to evaluate whether the periflagellar pocket location of TbPIP39 was an artefact of indirect immunofluorescence we generated cell lines with TbPIP39 fused to mNeon (Figure S1A) and used live cell imaging to detect the fluorescent protein in stumpy forms restrained in a hydrogel matrix. Figure S1B (movie S1A) revealed that in stumpy forms the fluorescent protein localised at the same periflagellar pocket site as previously visualised by immunofluorescence microscopy, a profile also seen with a YFP-TbPIP39 fusion protein (shown later in Figure 7C). Moreover, the signal redistributed to glycosomes in parasites exposed to *cis*-aconitate for 2 h, and 24 h (movie S1B, movie S1C).

Combined our results demonstrated that TbPIP39 was positioned close to, but not in, the flagellar pocket of the stumpy form parasites and distributed to glycosomes within 1-2h of the initiation of their differentiation to procyclic forms.

### Glycosomal dynamics during differentiation upon TbPIP39 depletion

During differentiation between stumpy forms and procyclic forms there is turnover of the glycosomal population presumably contributing to the metabolic adaptation of the parasites as they enter the tsetse fly. To determine whether TbPIP39 recruitment contributed to the control of glycosomal turnover and maturation during differentiation, we analysed the distribution of glycosomal aldolase and the lysosomal marker p67 during differentiation with TbPIP39 either depleted or not by RNAi. Previously, lysosomal and glycosomal staining patterns during differentiation have been subjectively categorised according to the distribution of aldolase and p67 in several cytological subtypes (A-E)(23). To determine if glycosomal/lysosomal dynamics were perturbed by reduced TbPIP39 recruitment during differentiation, pleomorphic *T. brucei* EATRO 1125 AnTat1.1. 90:13 bloodstream form parasites – competent for stumpy formation and inducible RNA interference – were generated able to deplete TbPIP39 under doxycycline regulation. When grown in mice provided either with or without doxycycline in their drinking water, uniform populations of stumpy forms were generated where TbPIP39 mRNA was targeted for RNAi, or not. In mice, the low levels of TbPIP39 were not significantly further reduced by RNAi and normal differentiation to stumpy forms occurred, as previously seen (18). However, when these stumpy form populations were induced to differentiate to procyclic forms with *cis*-aconitate, the normal increase of TbPIP39 levels was not observed over 24h, supporting RNAi mediated depletion during differentiation (Figure 6A). Over this period, procyclin expression occurred but was less efficient than in uninduced parasites, highlighting delayed or reduced differentiation with a reduction in TbPIP39, as previously reported (18).

**Figure 6.**
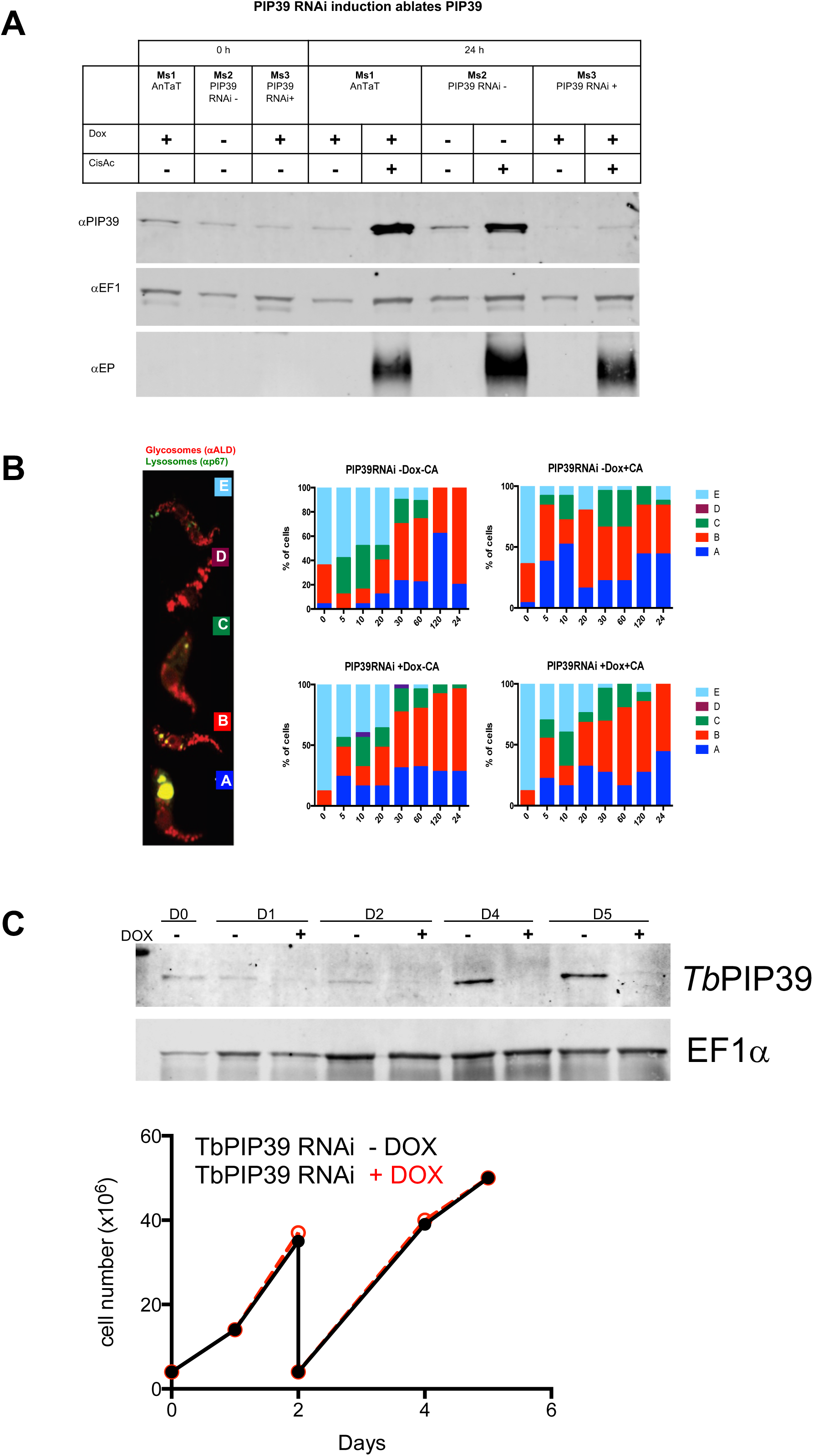
A. RNAi depletion of TbPIP39. *T. brucei* EATRO 1125 AnTat1.1 TbPIP39 RNAi cells or *T. brucei* EATRO 1125 AnTat1.1 cells were grown in mice, with or without doxycycline induction. The generated stumpy cells were then exposed, or not, to *cis*-aconitate to initiate differentiation (with doxycyline remaining in the RNAi-induced samples) and TbPIP39 protein detected at 24h. TbPIP39 levels increase during differentiation (ms1, ms2) but this is greatly reduced in induced RNAi samples (ms3). EF1 alpha provides a loading control, this being a little higher in cells exposed to *cis*-aconitate due to their replication as differentiated procyclic forms. EP procyclin staining shows relative differentiation in the presence or absence of *cis*-aconitate.
B. Distribution of the glycosomal marker aldolase and the lysosomal marker p67 during differentiation between stumpy and procyclic forms. The panel shows the prevalence of different categories of lysosomal/glycosomal staining (as defined by Herman et al., 2008 (Reference [23]), and shown at the left-hand side) at time through differentiation when TbPIP39 was depleted by RNAi or not.
C. Western blot of TbPIP39 in cells differentiated to proliferative procyclic forms with TbPIP39 RNAi induced or not. EF1alpha shows the loading control. The lower panel shows that by day 4 and 5 the cells remain proliferative (the cells were passaged at day 2) despite expressing significantly less TbPIP39.

To determine the consequences of TbPIP39 RNAi for the glycosomal and lysosomal configurations, cells were assayed during differentiation for the abundance of each glycosomal/lysosomal category (A to E) subjectively defined by (23) (Figure 6B). We observed no clear change in the distribution of the glycosome and lysosomal signal either in the presence or absence of *cis*-aconitate, or when TbPIP39 was depleted or not. The differentiated cells were also allowed to proliferate as procyclic forms for five days after the initiation of differentiation with TbPIP39 RNAi depletion maintained or not with doxycycline. Figure 6C demonstrates that although TbPIP39 remained significantly reduced with RNAi induction in the differentiated procyclic cells, these grew at an equivalent level to cells where TbPIP39 was not depleted.

We conclude that preventing the accumulation of TbPIP39 by RNAi does not significantly alter the changes in the glycosomal dynamics during differentiation from stumpy forms to procyclic forms. Moreover, differentiated procyclic forms do not require abundant TbPIP39 to sustain *in vitro* growth. Instead the dominant function of TbPIP39 under the growth conditions used appears to be restricted to regulating the efficiency of differentiation between bloodstream and procyclic forms.

### TbPTP1 and REG9. 1 localise close to TbPIP39 in stumpy forms

TbPIP39 is negatively regulated by the tyrosine phosphatase TbPTP1 that acts as an inhibitor of differentiation in stumpy forms. Previously, TbPTP1 had been difficult to localise using antibody specific for that molecule (16), and so we clarified its localisation with respect to TbPIP39 using an inducible N-terminally Ty1 epitope tagged copy (Figure 7A). Here, TbPTP1 was detected at the discrete periflagellar pocket location, with TbPIP39 co-labelling confirming that the location of the two signalling phosphatases was coincident in stumpy forms (Figure 7B). After 1h exposure to *cis*-aconitate, TbPIP39 relocated to glycosomes as seen earlier, whereas TbPTP1 became more diffuse at the periflagellar pocket site and by 4 h the signal was distributed throughout the cell body in the differentiating cells.

**Figure 7.**
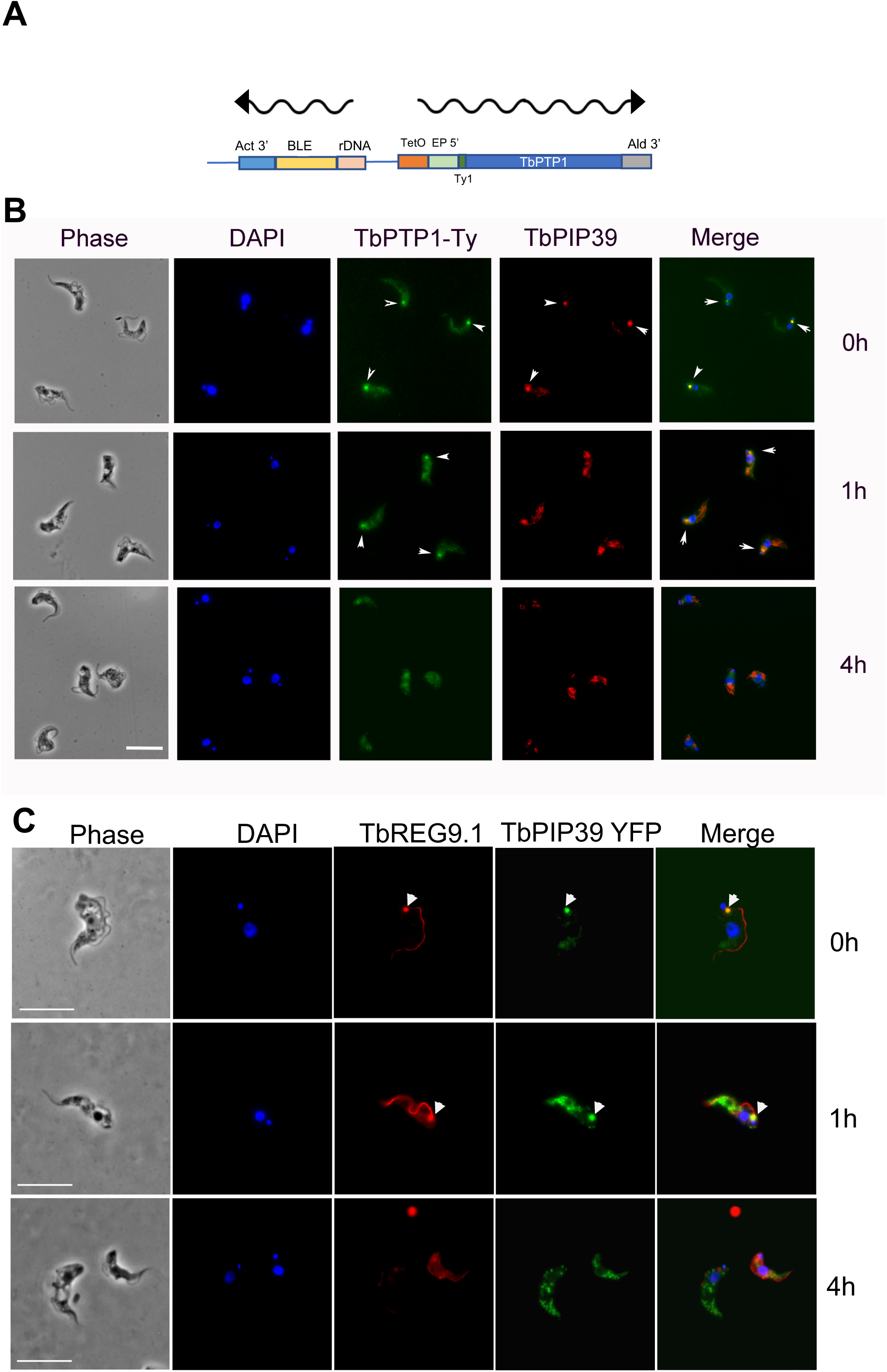
A. Schematic representation of the construct used to N-terminally tag TbPTP1 with the Ty1 epitope tag.
B. Co-localisation of TbPTP1-ty (green), and TbPIP39 (red) in stumpy forms (0h) and after 1h and 4h exposure to *cis*-aconitate. The TbPTP1 signal is detected as an ectopically expressed copy incorporating an N terminal Ty1 epitope tag to allow its detection. The TbPIP39 signal is detected using an antibody recognising that protein. The cell nucleus and kinetoplast are labelled with DAPI (blue). TbPTP1 colocalised with TbPIP39 in stumpy forms (arrowheads) but after 1h, TbPIP39 has relocated to glycosomes, while TbPTP1 has become more diffuse, albeit some remaining concentrated in the periflagellar pocket region (arrowheads). At 4h, TbPIP39 is glycosomal while TbPTP1 is diffuse throughout the cell body. Bar=10µm.
C. Detection of YFP tagged TbPIP39 (green) in stumpy form cells and its location with respect to REG9.1 (red). DAPI (blue) denotes the position of the cell nucleus and kinetoplast. Bar=15µm.

The TbPIP39/TbPTP1 node was also similar to the location of REG9.1, a regulator of stumpy specific transcripts previously observed at the periflagellar pocket region of stumpy forms, in addition to along the flagellum or FAZ (26). Therefore, we colocalised REG9.1 with TbPIP39 using a REG9.1 specific antibody and YFP-tagged TbPIP39 during differentiation and observed redistribution form the periflagellar pocket node to the cytoplasmic distribution previously in stumpy forms within 4 h (Figure 7C). Hence, TbPTP39, TbPTP1 and REG9.1 are all colocalised in stumpy forms at the same cellular site before their redistribution and separation at the onset of differentiation.

### The periflagellar location of TbPIP39 and TbPTP1 coassociates with flagellar pocket ER

The observed location of TbPIP39 in stumpy forms was reminiscent of a specialised region of the ER at the flagellar pocket identified by electron tomography of procyclic form cells (22). This region of the cell is defined by the presence of TbVAP, a flagellum attachment zone (FAZ) ER membrane contact protein. TbVAP depletion by RNAi in procyclic forms results in loss of the FAZ ER and flagellar pocket ER, but no detectable growth deficit (22). To determine if the TbPIP39 location in stumpy forms associated at the flagellar pocket ER, we colocalised TbPIP39 with TbVAP detected by its incorporation of N terminal mNEON Ty1 epitope tag and expression in pleomorphic cells. Figure 8A demonstrates that the TbPIP39 and TbVAP signals are closely-located in the flagellar pocket region of stumpy forms, but not completely coincident, with TbPIP39 sometimes between two separated TbVAP foci. Matching previous analysis in procyclic form cells (22), TbVAP signal also extended along the flagellum attachment zone with further staining in the cell body.

**Figure 8.**
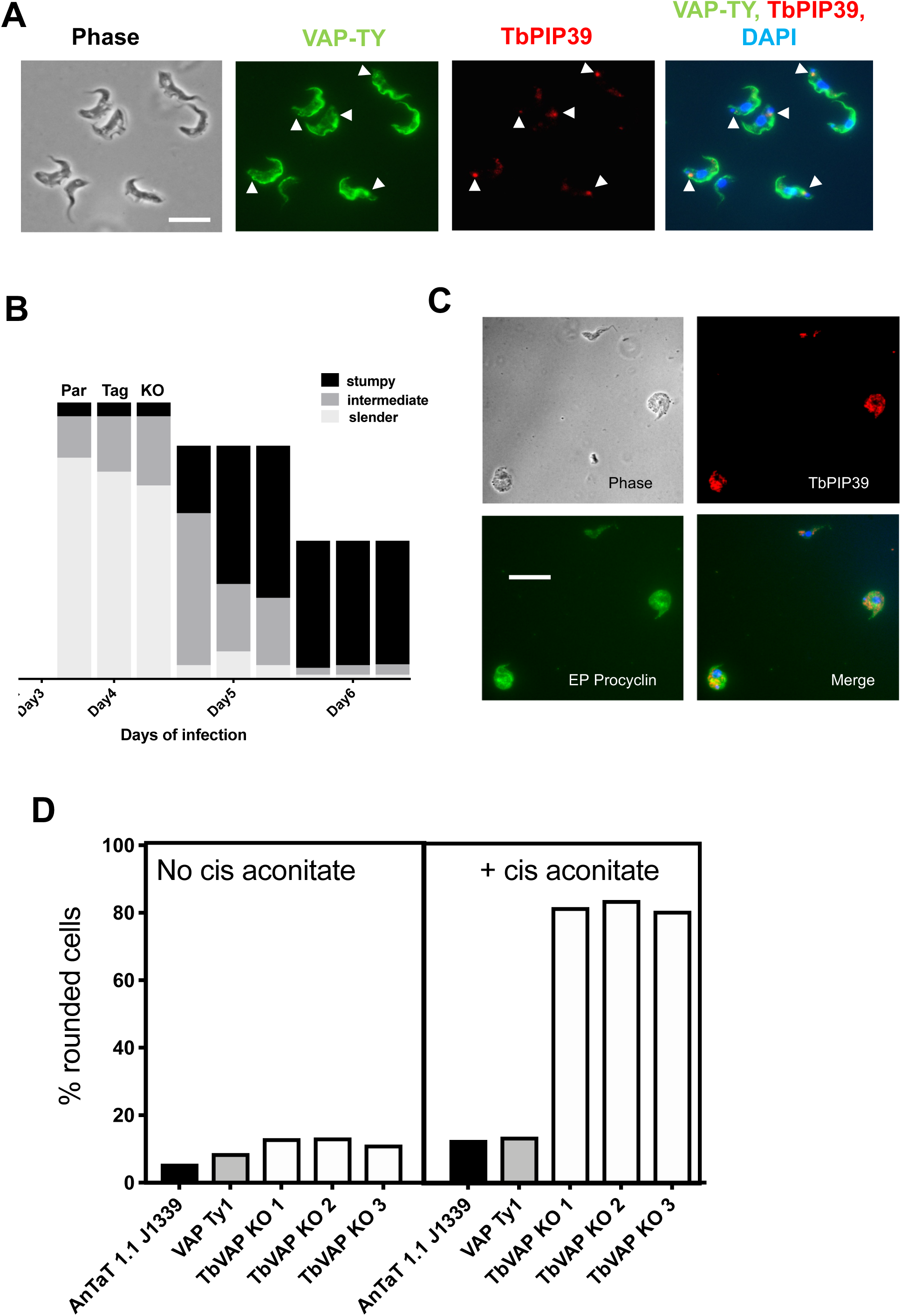
A. Detection of Ty1 epitope tagged TbVAP in stumpy form cells and its location with respect to TbPIP39. The Ty1 tagged TbVAP (green) is detected at the flagellar pocket region, along the flagellum attachment zone and in the cell body. TbPIP39 (red) is closely proximal at the flagellar pocket region but not precisely coincident. DAPI (blue) denotes the position of the cell nucleus and kinetoplast. Arrowheads indicate the region of TbPIP39. Bar=15µm.
B. Proportion of slender, intermediate and stumpy forms in TbVAP single knock out cells versus parental cells, or epitope tagged TbVAP cells.
C. TbVAP single allele replacement mutants 24h after the initiation of differentiation to procyclic forms. The differentiating cells (indicated by their expression of EP procyclin, green) are swollen and lose integrity. TbPIP39 is punctate throughout the cell, indicating a glycosomal rather than periflagellar pocket distribution. The merged panel shows TbPIP39 (red), EP procyclin (green) and DAPI (blue) revealing the nucleus and kinetoplast. Bar=25µm.
D. Quantitation of cells which exhibit a rounded morphology. Parental *T. brucei* AnTat1.1 J1339 cells, TbVAP1-mNeon-Ty1 expressing cells or three TbVAP single allele deletion mutants are shown. In each case the percentage of rounded cells is shown after 24h either in the absence or presence of *cis*-aconitate as a differentiation stimulus. At least 250 cells were scored for each cell line and each condition.

To explore the association of TbPIP39 and TbVAP at this periflagellar pocket location in more detail, we exploited CRISPR to delete the TbVAP gene in pleomorphic bloodstream form trypanosomes (24). Thus, the Cas9 expressing cell line *T. brucei* EATRO 1125 AnTat90:13 J1339 (25) was transfected with repair templates targeted to the TbVAP gene genomic location, and mutants were isolated (Figure S2A). The resulting cell lines were found to retain a TbVAP gene copy despite their successful insertion of the drug resistance gene (Figure S2B); further attempts to delete the remaining allele were unsuccessful. These TbVAP depleted cells exhibited stumpy formation *in vivo* with similar kinetics as wild type cells (Figure 8B) allowing the location of TbPIP39 and its relocation after the initiation of differentiation to procyclic forms to be assayed.

In TbVAP single KO stumpy forms, TbPIP39 was detected at the periflagellar pocket location, similar to that seen in wild type parasites. Correspondingly, when cells were induced to differentiate to procyclic forms with *cis*-aconitate, TbPIP39 relocated to glycosomes and the cells expressed procyclin, indicating that TbVAP depletion did not alter glycosomal TbPIP39 loading or differentiation (not shown). However, when the TbVAP single KO cells were analysed after 24h in differentiation conditions (*i.e.* with *cis*-aconitate) the parasites appeared enlarged and rounded and were dying, unlike wild type parasites at this time point (Figure 8C) (mean=82% from an analysis of three independent null mutant lines, versus 13% and 14% in the parental and VAP-Ty1 line, respectively, n=250 cells per sample; Figure 8D). Thus, both alleles of TbVAP are necessary for the viability of differentiating cells, but not for the normal relocation of TbPIP39 early in the process.

## Discussion

The differentiation of African trypanosomes between life cycle stages is enacted rapidly upon transition from the blood of mammalian hosts to the midgut of the tsetse fly. We have shown previously that two phosphatases are important in the signalling of the changes from one environment to the next, TbPTP1 and TbPIP39. Of these TbPIP39 is glycosomal in procyclic forms, signalled through its C terminal PTS 1. Here we show that when first made in stumpy forms, TbPIP39 is not glycosomal but rather is localised at a periflagellar pocket region of the parasite where it colocalises with the differentiation inhibitor, TbPTP1. However, upon reception of the differentiation signal TbPIP39 is rapidly (within approximately 20 minutes) relocated into glycosomes, whereas TbPTP1 becomes dispersed to a non-glycosomal, possibly cytosolic site. It is also coincident with the location of a regulator of stumpy form transcripts, REG9.1. Interestingly, this periflagellar pocket site is close to the specialised FAZ endoplasmic reticulum, defined by TbVAP, that may represent a site of glycosomal biogenesis in the differentiating cells. We propose this molecular node comprising TbPIP39, TbPTP1 and REG9.1 generates a ‘stumpy regulatory nexus’ (STuRN) where the events initiating differentiation between bloodstream and procyclic forms occur.

The biogenesis of peroxisomes and glycosomes can involve new organelles that arise from pre-existing peroxisomal/glycosomal structures or *de novo* synthesis from the endoplasmic reticulum. However, in eukaryotes subject to environmental change, peroxisome composition can be modulated to allow metabolic adaptation, and this can be achieved by *de novo* loading from the ER (27) as well as the growth and division of existing glycosomes. The dynamics of environmental adaptation in procyclic form parasites have been analysed both during differentiation and upon exposure of procyclic forms to low and high glucose concentrations. In differentiation, autophagy is considered important due to the co-association of lysosomes and glycosomes early in the transition between stumpy and procyclic forms (23). The assessment of this is relatively subjective and we observed limited co-association between the lysosomal marker p67 and glycosomal aldolase during synchronous differentiation, at least in the first 2 hours. In contrast, TbPIP39 relocated from the STuRN to glycosomes within 20 minutes, this region being neither the flagellar pocket lumen or the flagellar pocket membrane. Instead the STuRN was adjacent to a specialised region of the endoplasmic reticulum that has been visualised by electron tomography of procyclic forms cells (22) and is also the site of ER concentration in procyclic forms in low glucose. In this region of the cell, ER is associated with both the flagellar pocket and flagellum attachment zone, the region being defined by TbVAP, an orthologue of VAMP associated protein. This molecule is proposed to co-ordinate ER in this region of the cell through linking the specialised four microtubules positioned at the flagellar pocket region of the cytoskeleton to the endoplasmic reticulum or controlling interaction between central ER and the flagellar pocket associated ER. In a previous study, procyclic form viability was not compromised by efficient RNA interference targeting TbVAP suggesting that it is not essential or that significant depletion of its protein levels after RNA interference does not compromise cell viability and replication. By CRISPR/Cas9 mediated gene deletion we also found that bloodstream forms were viable after the deletion of one allele although both alleles could not be deleted. This may reflect that the protein is essential, contrasting with RNAi-depletion experiments, although technical reasons cannot be excluded. Interestingly, however the TbVAP single KO mutants – although able to initiate differentiation between stumpy and procyclic forms – lost cell integrity after 24h, and appeared swollen and balloon like. This suggests that the levels of this molecule are important during differentiation.

Our data invoke a model where in stumpy forms TbPIP39 is poised for glycosome recruitment through its recruitment to the STuRN in a pre-glycosomal concentration similar to pre-peroxisomal vesicles (Figure 9). At this site, the presence of TbPTP1 inhibits its activity. With the initiation of differentiation however, TbPTP1 is inactivated and the TbPIP39 protein is activated by phosphorylation through the activity of an -as yet-unidentified kinase and assembled into glycosomes where it can no longer be accessed by TbPTP1. Removing the inhibitor TbPTP1 from its substrate, TbPIP39, renders the differentiation signalling irreversible – a commitment event that has been mapped to approximately 1 hour after exposure to citrate/*cis*-aconitate (18, 28), coincident with the dispersal of TbPIP39 and TbPTP1 in differentiating cells. The glycosomal pool rapidly turns over through autophagy during differentiation, but the loading of new glycosomes generated at the flagellar pocket region allows the remodelling of the glycosomal pool to its procyclic composition. It remains to be established whether TbPIP39 activity is necessary in the mature glycosomes that arise during differentiation or in procyclic forms, since TbPIP39 RNAi did not affect the growth of differentiated cells *in vitro*. It will be interesting to explore the fitness of procyclic forms depleted of TbPIP39 in different culture conditions more closely mimicking conditions in the fly gut, such as when alternative carbon sources are available (29, 30).

**Figure 9.**
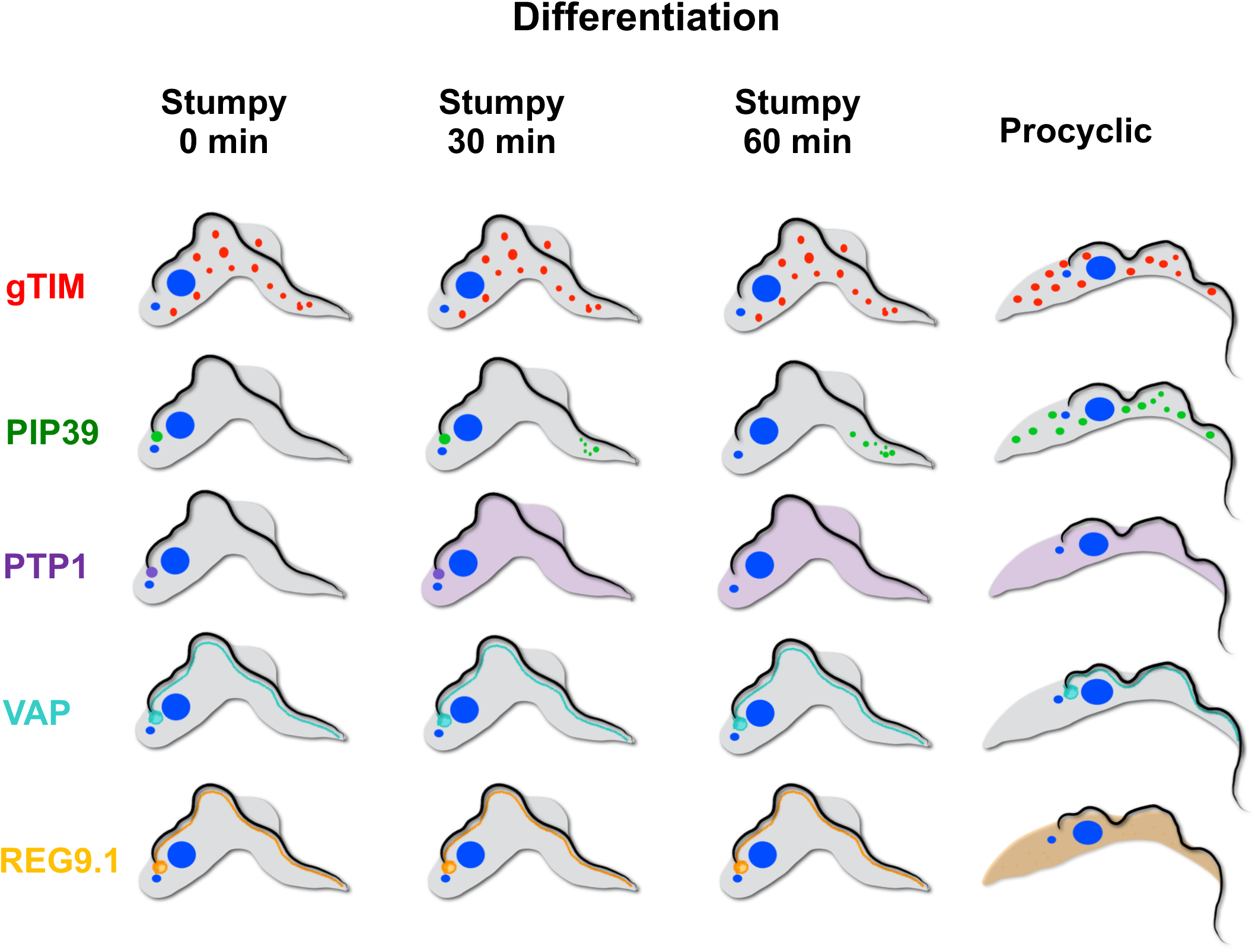
Schematic representation of the distribution of gTIM (representing glycosomes) TbPIP39, TbPTP1, TbVAP and REG9.1 in stumpy forms and at 30 minutes and 60 minutes after exposure to *cis*-aconitate and in procyclic forms.

The synchronous differentiation of trypanosome parasites and their developmental adaptation of glycosomal composition provides a unique capability to explore the dynamics of organellar development in an evolutionary divergent eukaryotic model. This study provides the first temporal and positional tracking of signalling molecules, organellar compartments and contact points at the STuRN, a potential site of glycosomal biogenesis/regeneration as the parasite initiates its metabolic adaptation to a nutritionally distinct environment. Coupled with the high definition understanding of the structural organisation and cytoskeletal interactions in this region of the highly-ordered parasite cell (31–33), trypanosomes provide an invaluable model for the precise regulation and kinetics of inter-organellar exchange in a eukaryotic cell.

## Acknowledgements

We thank Professor Paul Michels, University of Edinburgh, for advice and constructive comments on the manuscript. DVS was supported by Erasmus+Mobility grant. DRR was supported by internal grants (CNRS and university of Bordeaux) and support from the ANR, LABEX ParaFrap, ANR-11-LABX-0024 (http://www.labex-parafrap.fr/fr). ME is supported by DFG grants EN305, GRK2157 and SPP1726. Research in KRM’s laboratory is funded by a Wellcome Trust Investigator award (103740/Z14/Z) and Royal SocietyWolfson Research Merit award (WM140045).

The funders had no role in study design, data collection and interpretation, or the decision to submit the work for publication.

## Materials and Methods

### Parasites

#### Cell lines and culturing in vitro

Pleomorphic *Trypanosoma brucei* EATRO 1125 AnTat1.1 90:13 (TETR T7POL NEO HYG) (19) and EATRO AnTat 1.1 J1139 (25) parasites were used throughout.

Pleomorphic bloodstream and double marker 29-13 procyclic form trypanosomes (34) were cultured *in vitro* in HMI-9 (35) medium at 37°C 5% CO2 or in SDM-79 (36) medium at 27°C respectively.

The following selective drugs were used: Hygromycin (2.5 μG/mL), Puromycin (0.5 μG/mL) and Blasticidin (2.5 μG/mL).

#### In vivo studies

Trypanosome infections were carried out in female healthy outbred MF1 mice at least 10 weeks old, immunocompromised with 25mg/ml cyclophosphamide delivered intraperitoneally 24 h prior to trypanosome infection.

No blinding was performed and the animals were not subject to previous procedures or drug treatment. Animal experiments were carried out according to the United Kingdom Animals (Scientific Procedures) Act under a license (PPL60/4373) issued by the United Kingdom Home Office and approved by the University of Edinburgh local ethics committee. Animals were kept in cages containing 1-5 mice on a 12h daylight cycle and maintained at room temperature.

*In vivo* growth involved intraperitoneal injection of 10^5^ parasites into cyclophosphamide-treated mice and the course of parasitaemia was recorded by performing daily tail-snips to estimate parasite numbers using a ‘rapid matching’ method involving visual comparisons of live parasites in blood by microscopy with a published standardized chart of parasite numbers per ml (37).

Ectopically expressed gene expression was induced by inclusion of doxycycline (200 mg/mL in 5% sucrose) in the drinking water, with control mice being provided with 5% sucrose alone. Between 2 and 3 mice were used per group.

Stumpy-enriched populations were obtained 6–7 d after infection by DEAE cellulose purification (38).

For the initiation of differentiation conditions were used as described in (17).

#### Parasite transfection

Parasite transfection was by Amaxa nucleofection according to previous detailed methods for pleomorphic (39) and 29-13 procyclic form parasites (40).

### Plasmid construction and cell line generation

#### Generating endogenously tagged TbPIP39 pleomorph cell lines

Primers 1.-4. were used to endogenously tag the N-terminus of TbPIP39 using pPOTv4YFP and pPOTv6 mNEONgreen plasmids according to (41).

#### Generating endogenously tagged TbPTP1 pleomorph cell lines

The TbPTP1 open reading frame was amplified from *T. brucei* EATRO 1125 AnTat1.1 wild-type genomic DNA by PCR using Primer 5 and Primer 6 (Table S1) with SpeI and Bgl II restriction sites for insertion into the pDex577-Y vector (42) for tetracycline-inducible overexpression with an N-terminal Ty1 epitope tag. The resulting overexpression constructs were linearized with NotI and transfected into *Trypanosoma bucei* EATRO 1125 AnTat1.1 90:13 pleomorphs cells. Several independent cell lines were isolated and their growth analyzed *in vitro* or *in vivo* in the presence or absence of tetracycline, or doxycycline, respectively. Expression was confirmed by Western blotting using an anti-Ty1 antibody.

#### Generating endogenously tagged and knockout TbVAP pleomorph cell lines

The LeishGEdit program was used (24) to design oligonucleotide primers (Table S1 Primers 7-11) to produce DNA fragments and sgRNAs for the production of Ty1 mNEONGreen tagged VAP and KO VAP cell lines. To create the KO and endogenously tagged pleomorph cell lines EATRO AnTat 1.1 J1139 cells were transfected as described in(25).

Several independent cell lines were isolated and their growth analyzed *in vitro* or *in vivo*. Pleomorph cell lines with the mNEON-Ty1 tagged TbVAP was identified by Western blotting and immunofluorescence using an anti-Ty1 antibody.

Several TbVAP knockout cell line candidates were isolated, and genomic DNA were purified (QIAGene GenomicDNA kit). The genomic DNAs were used in PCR reactions to confirm the presence of the Blastocidine drug resistance cassette (replacing the endogenous TbVAP) (Table S1 primers 7-10) and also the lack of the endogenous TbVAP gene (Table S1 primers 14-15). TbPIP39 RNAi lines were described in (17).

### Western blotting, immunofluorescence and confocal, Immuno Electronmicroscopy and live cell microscopy

Protein expression analyses by Western blotting were carried out according to (17) with antibody concentrations detailed in Table S2.

Approximately 1×10^9^ EATRO AnTat1.1 90:13 stumpy (around 90-95% cells of the isolated cells were stumpy forms, the rest were intermediates) cells from mice were purified on DE52 column in PSG buffer. After purification, cells were resuspended in HMI9 at 4×10^6^/ml and left in a 37°C CO_2_ incubator for 60 mins to recover. Then the culture was divided into two aliquots and ∼5×10^8^ stumpy cells treated with *cis*-aconitate (+ CA sample) and 5×10^8^ stumpy cells left untreated (-CA sample). After 60 mins CA induction cells were harvested by centrifugation, washed in 15 ml of vPBS and repelleted, before resuspension in 10 ml vPBS (2.5 ×10^8^ cells/10 ml). Immunofluorescence was carried out according to (20) using the antibody concentrations in Table S2. Phase–contrast and Immunofluorescence microscopy images were captured on a Zeiss axioskop2 (Carl Zeiss microimaging) with a Prior Lumen 200 light source using a QImaging Retiga 2000RCCD camera; the objective was a Plan Neofluar ×63 (1.25 NA). Images were captured via QImage (QImaging). Cells were captured for confocal microscopy or processed for immunoelectron or live cell microscopy as described in Text S1.

## Text S1

### Supplementary material and methods

#### Confocal imaging

Confocal imaging used a Leica SP5 confocal laser scanning microscope, using 63x oil immersion objective (NA = 1.4) and 4.2x digital zoom. The green channel was imaged using a 488-nm argon laser, and the red channel was imaged using a 543-nm helium/neon laser. The final image was acquired using Volocity Software (www.perkinelmer.co.uk) version 4.4. 3D fluorescence microscopy was performed using a fully automated Leica DMI6000B inverse microscope equipped with a 100x (NA 1.4) oil immersion objective and a Leica DFC365 FX CCD camera. Fluorescent and differential interference contrast (DIC) image stacks were acquired by recording stacks of 50 images with a step size of 137 nm. Fluorescent image stacks were deconvolved using Huygens Essential (SVI, Hilversum, Netherlands). Images were visualised as z-stack projections using Fiji(43), or as volumetric representations using Amira (Thermo Fisher Scientific).

#### Electron microscopy

After incubation of cells in an equal volume of vPBS and 8% paraformaldehyde, cells were pelleted at 1,000x g for 15 min at 4°C. The supernatant was removed and the pellet dehydrated with 30% ethanol at 4°C for 30min. The pellet was then taken through the following series of methanol dehydrations, 60%, 90%, 3 × absolute (30 min each) at −20°C. After dehydration, the pellet was incubated in 2mL of 30% HM20 Lowicryl monostep EM resin (EMS – 14345) in methanol at −20°C, 30min, then taken through increasing concentrations of HM20 (60 min each) then 3 × 100% absolute HM20 at −20°C. The pellet was polymerised at −20°C for 24hr using UV light at 360nm, then left at −20 °C for 24h. The block was cut using a diamond knife and 100nm thin sections were incubated on 100µL droplets of 100mM glycine in PBS 2 × 10 min. Sections were then probed with primary antibody 25µL droplets of rabbit anti-PIP39 diluted 1:100 in incubation buffer (PBS, containing 0.1% Tween 20, 0.1% BSA, pH 7.3), for 2h at room temp. After washing, 4 × 10 min in incubation buffer, grids were incubated in a 1:1 mix of protein A (EMS – 25285), plus protein G (EMS – 25315) 10 nm gold, diluted 1:25 in incubation buffer for 2h at room temp. Grids were washed 2 × 10 min in PBS, containing 0.1% Tween 20, 0.1% BSA, pH 7.3, then 2 × 10 min in PBS, pH 7.3, then 2 × 10 min in Milli-Q water. Grids were stained in 2% uranyl acetate for 1min at room temp then washed 4 × 10 min in Milli-Q water and visualised on a Technai 12 at 120Kv.

#### Live cell microscopy

##### Sample preparation for live cell microscopy

For studying stumpy to procyclic form differentiation by live cell microscopy 25 µL of pH7.6, 0.6 M, filtered *cis*-aconitate was added to 2.5 ml of stumpy culture (4×10^6^/ml) to induce differentiation.

After 10 mins 1mL sample of the induced culture was taken to concentrate for the hydrogel fixation. Samples were spun for 10 min at 1400 rpm at 37°C and 980 µL supernatants were removed after the spin cells were carefully resuspended in 20 µL culture media.

Immobilisation of living cells was performed with a two component hydrogel, consisting of 8-arm poly ethylene glycol (PEG) norborene and linear PEG-dithiol, essentially as described in (44). 4µl of hydrogel produced with TDB was mixed with a 2 µL solution containing 4×10^5^ differentiating stumpy cells and incubated 1 minute on ice. The mixture was transferred to a glass bottom cell culture dish (FluoroDish, FD35, World Precision Instruments) and allowed to polymerise at room temperature. Cells were viable in the hydrogel for at least 60 minutes (Glogger et al., 2017). In the experiments for this work, the hydrogel concentration was chosen so as to prevent translocation of cells, but still allow flagellar movement. As a control, 100 µL of the induced culture was taken and spun at the same speed and time as the immobilised sample. The cell pellet was carefully resuspended in ml of warm TDB (2×10^5^/ml) and was used to overlay the hydrogel droplet containing *cis*-aconitate induced stumpy cells.

Live imaging was performed with a Leica DMI6000B fully automated inverse microscope equipped with a 100x (NA1.4) oil immersion objective and a pco.edge SCMOS camera (PCO, Kelheim, Germany). Time series were recorded with 100 fps (100 ms resolution), which allowed the accurate detection of dynamic fluorescent signals in the cytoplasm of the living cells. Continuous videos were recorded switching between the fluorescence excitation and DIC settings. Due to the repetitive, periodic beating of the flagellum and cell movement of the immobilised trypanosome, the fluorescent and DIC images of corresponding phase shifted time series could be overlaid using Fiji (Fig. S1B, 0h).

---

**Table S1.**
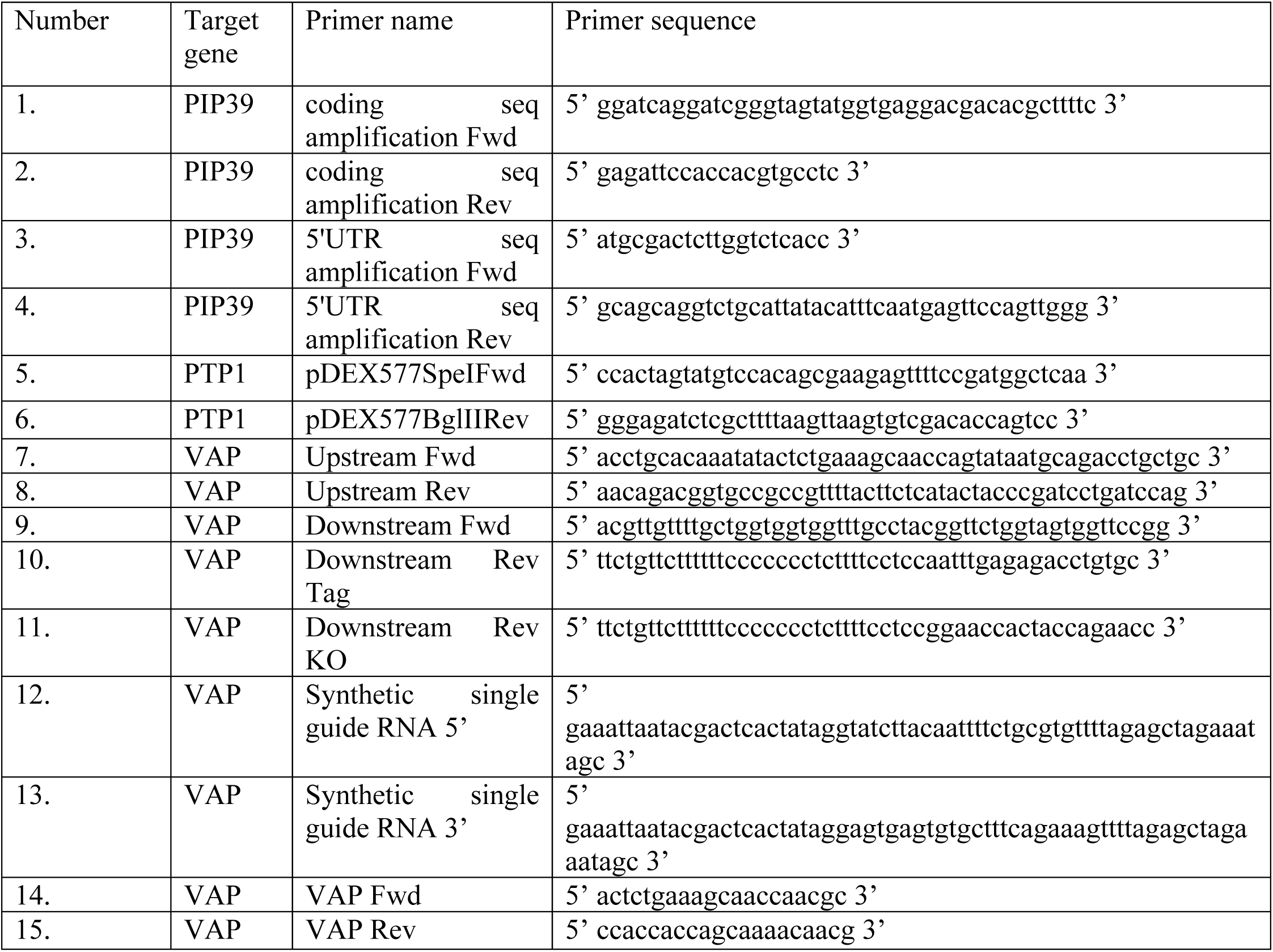
Oligonucleotides used in the study.

**Table S2:** Antibodies used in the study.

| Antibody name | Animal raised in | Dilution used on Western Blot (W)<br>or in immuno fluorescence assays (IF) |
| --- | --- | --- |
| PIP39 | Rb | 1:750 W, 1:300 IF |
| BB2 | Ms | 1:5 W, 1:4 IF |
| EF1 | Ms | 1:7000 W |
| EP procyclin | Ms | 1:3000 W, 1:250 IF |
| gTIM | Ms | 1:500 IF |
| Aldolase | Ms | 1:250 IF |
| p67 | Ms | 1:1000 IF |
| REG 9.1 | Rb | 1:1000 IF |
| Alexa568 Red | Ms/Rb | 1:500 IF |
| Alexa488 Green | Ms/Rb | 1:500 IF |
| Licor Red Rb | Ms/Rb | 1:7500 W |
| Licor Green Ms | Ms/Rb | 1:7500 W |

**Figure S1.**
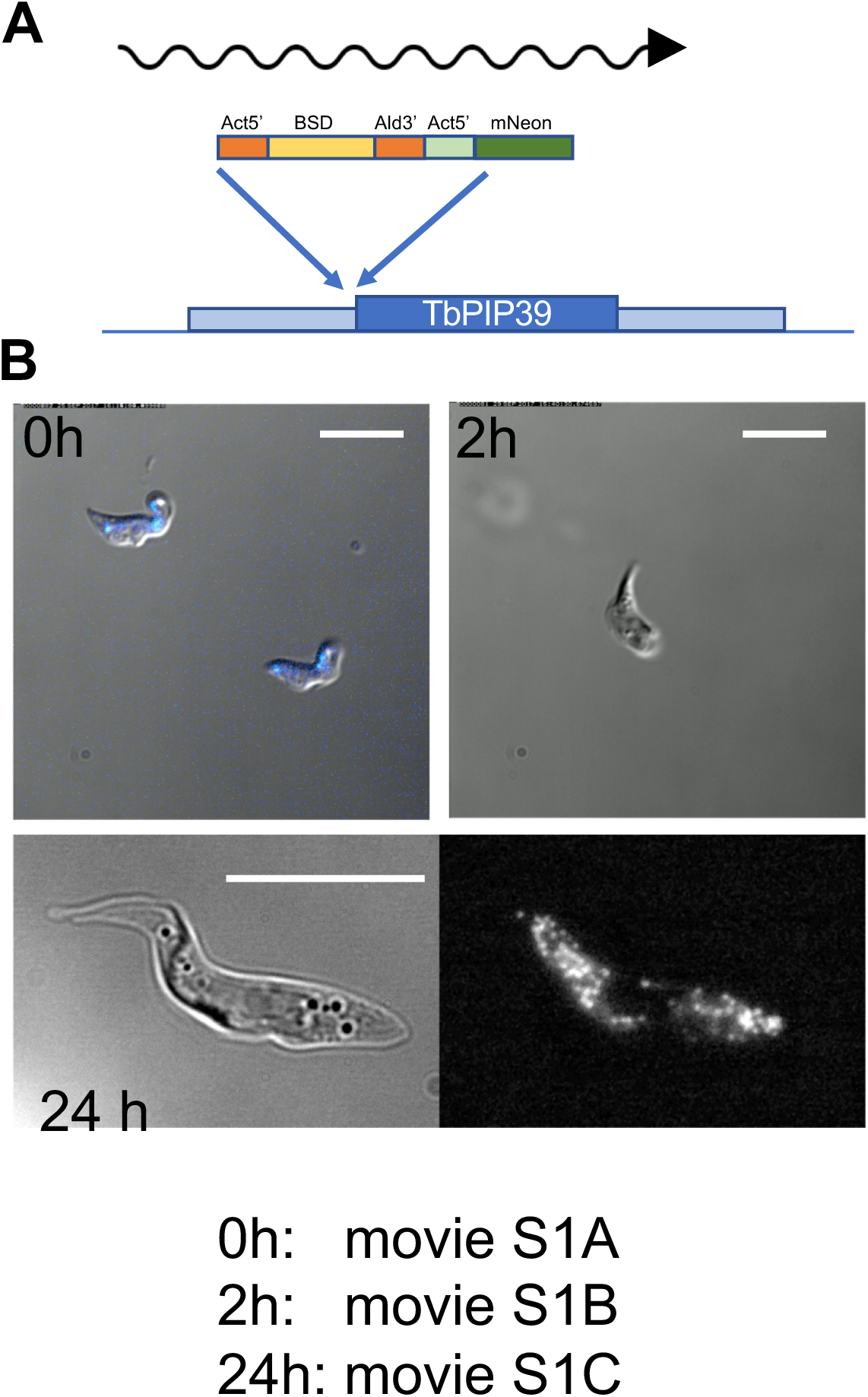
A. Schematic representation of the construct used to N-terminally tag TbPIP39 with mNeon.
B. Video images of TbPIP39 mNeon signal in stumpy forms (0h), or 2h or 24h after exposure to *cis*-aconitate to initiate differentiation. At 0h the signal is predominantly periflagellar pocket, though with some dispersed glycosomal signal. At 2h there is still some periflagellar pocket signal but stronger glycosomal signal. At 24h the signal is exclusively glycosomal. Bar=20µm.

**Figure S2.**
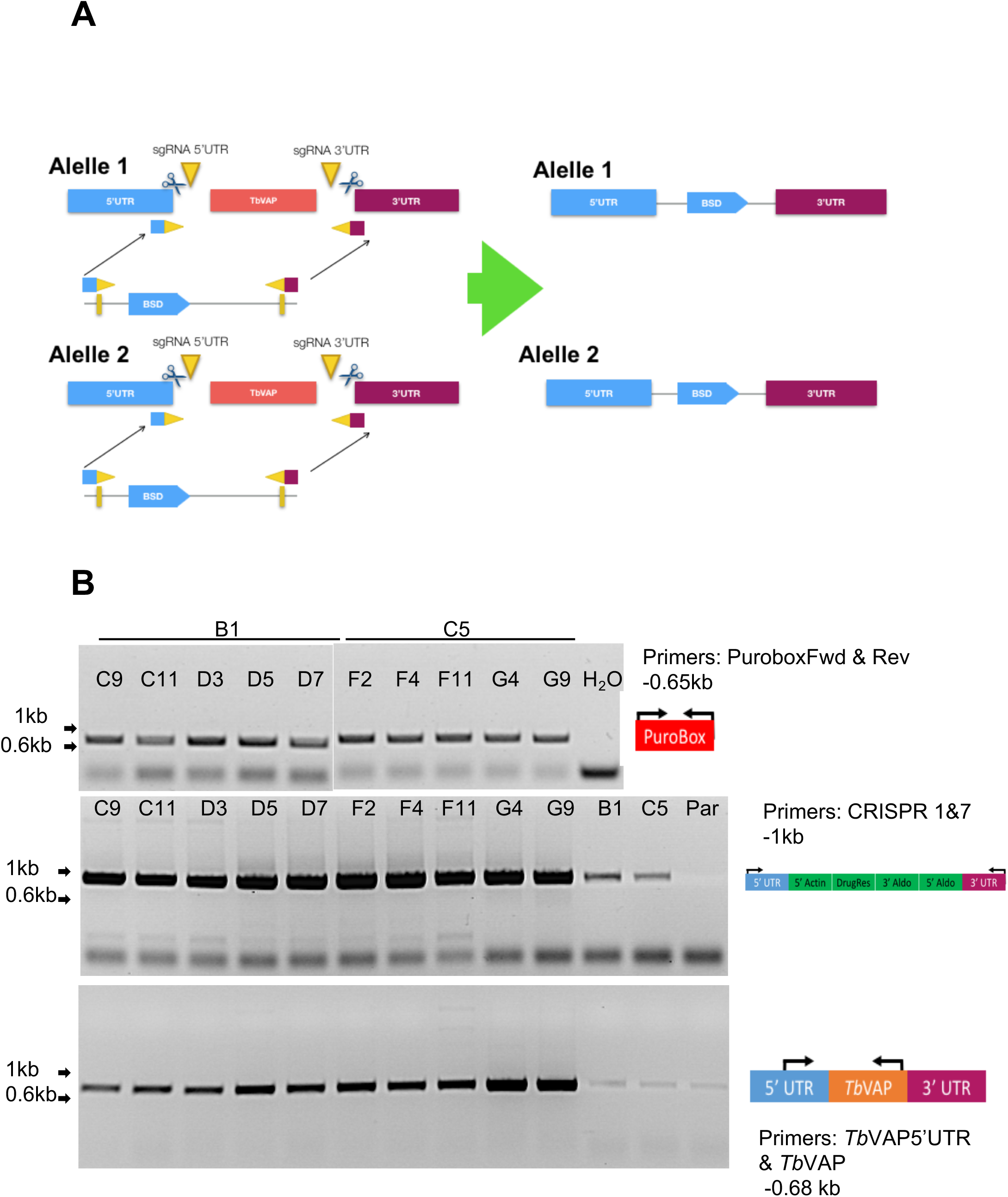
A. Schematic representation of the deletion of TbVAP alleles by CRISPR. The allele was deleted through integration of a blasticidin resistance cassette.
B. PCR analysis of the insertion of a TbVAP replacement construct using CRISPR. All cell lines derived have successfully incorporated the VAP replacement cassette but also retain an intact copy of the TbVAP gene.

